# External Globus Pallidus Arkypallidal Circuit Dynamics Gate Risk-Taking Behavior

**DOI:** 10.64898/2026.03.20.713182

**Authors:** David L. Haggerty, Beatriz D. Sorigotto, Armando G. Salinas, David M. Lovinger, Karina P. Abrahao

## Abstract

Exploration allows animals to gather information and adapt to changing conditions. Yet, it also exposes them to potential threats, requiring neural systems that weigh uncertainty and regulate behavioral transitions between cautious and exploratory states. These computations are distributed across cortical and subcortical networks, including the basal ganglia, which integrate sensory, motivational, and contextual information to shape action-selection behaviors. Within this circuitry, the globus pallidus externa (GPe) occupies a central but underappreciated role. Once viewed as a relay between striatum and downstream nuclei, the GPe is gaining recognition as a dynamic regulator that integrates diverse inputs and exerts bidirectional control over motor and cognitive processes. Here, we examine arkypallidal NPAS1-expressing GPe (GPe^NPAS1^) neurons, which form preferential inhibitory projections to the striatal matrix. Chemogenetic manipulations and *in vivo* calcium measurement reveal that GPe^NPAS1^ activity modulates and encodes risk-taking behavior sequences, identifying a circuit mechanism by which the GPe can regulate adaptive decision-making in risky contexts.

## Introduction

Adaptive behavior depends on an organism’s ability to explore its environment while avoiding danger. Exploration allows animals to gather information and adapt to changing conditions, but it also exposes them to potential threats, requiring neural systems that evaluate uncertainty and regulate transitions between cautious and exploratory states^1,2^. Such computations engage distributed cortical and subcortical networks that integrate sensory, motivational, and contextual information to guide action-selection^3–8^. The basal ganglia is central to this process, interfacing cortical representations of context and downstream motor systems to determine when to initiate, sustain, or suppress actions under varying degrees of uncertainty^2,3,9–11^.

Historically, the external globus pallidus (GPe) was viewed primarily as a relay between the striatum and downstream basal ganglia nuclei^12–16^. However, recent studies reveal that the GPe contains diverse, molecularly distinct neuron classes that form recurrent loops with the striatum and other basal ganglia structures^17–22^. Through these connections, the GPe may exert control over both motor output and higher-order cognitive processes, dynamically shaping behavioral flexibility. Despite this emerging view, how specific GPe neuron types encode and influence decisions remains poorly understood, and the circuit substrates that distinguish cognitive from motor control within the GPe are not well defined.

Here, we focus on NPAS1-expressing neurons of the GPe (GPe^NPAS1^), which send inhibitory gabaergic projections to the dorsal striatum that define the arkypallidal projection^23–25^. Using anatomical tracing and electrophysiology, we identify that GPe^NPAS1^ neurons preferentially innervate the striatal matrix rather than the striosomes - a pattern not previously described. Because the matrix is often associated with sensorimotor and associative processing, this projection positions GPe^NPAS1^ neurons to regulate the flexibility and sequencing of exploratory behaviors rather than their value^26^. Combining chemogenetic manipulation and *in vivo* fiber photometry, we show that GPe^NPAS1^ neurons modulate risk-taking behavior without altering gross locomotion, and that their activity encodes moment-to-moment transitions across the risk-taking spectrum. These results reveal a circuit mechanism through which the GPe shapes cognitive aspects of decision-making, providing a foundation for understanding how pallidal dynamics contribute not only to adaptive motor behavioral control, but also its associated cognitive processing.

### GPe^NPAS1^ neurons preferentially innervate the striatal matrix and exert stronger functional inhibition onto matrix SPNs

To determine whether GPe^NPAS1^ neurons differentially target striatal subcompartments, we bred Npas1-Cre-tdTomato with Nr4a1-eGFP mice to quantify GPe^NPAS1^ terminal expression in the matrix and striosome (Fig. 1a)^27^. Confocal imaging revealed dense Npas1⁺ terminal labeling throughout matrix regions in the dorsal striatum, whereas Npas1⁺ terminals were markedly sparse within striosome compartments (Fig. 1b–d). Quantification of terminal density, as a function of total labeling area within the regions of interest, confirmed a ∼4 fold enrichment of GPe^NPAS1^ terminals in the matrix relative to the striosomes (Fig. 1e).

**Figure 1.**
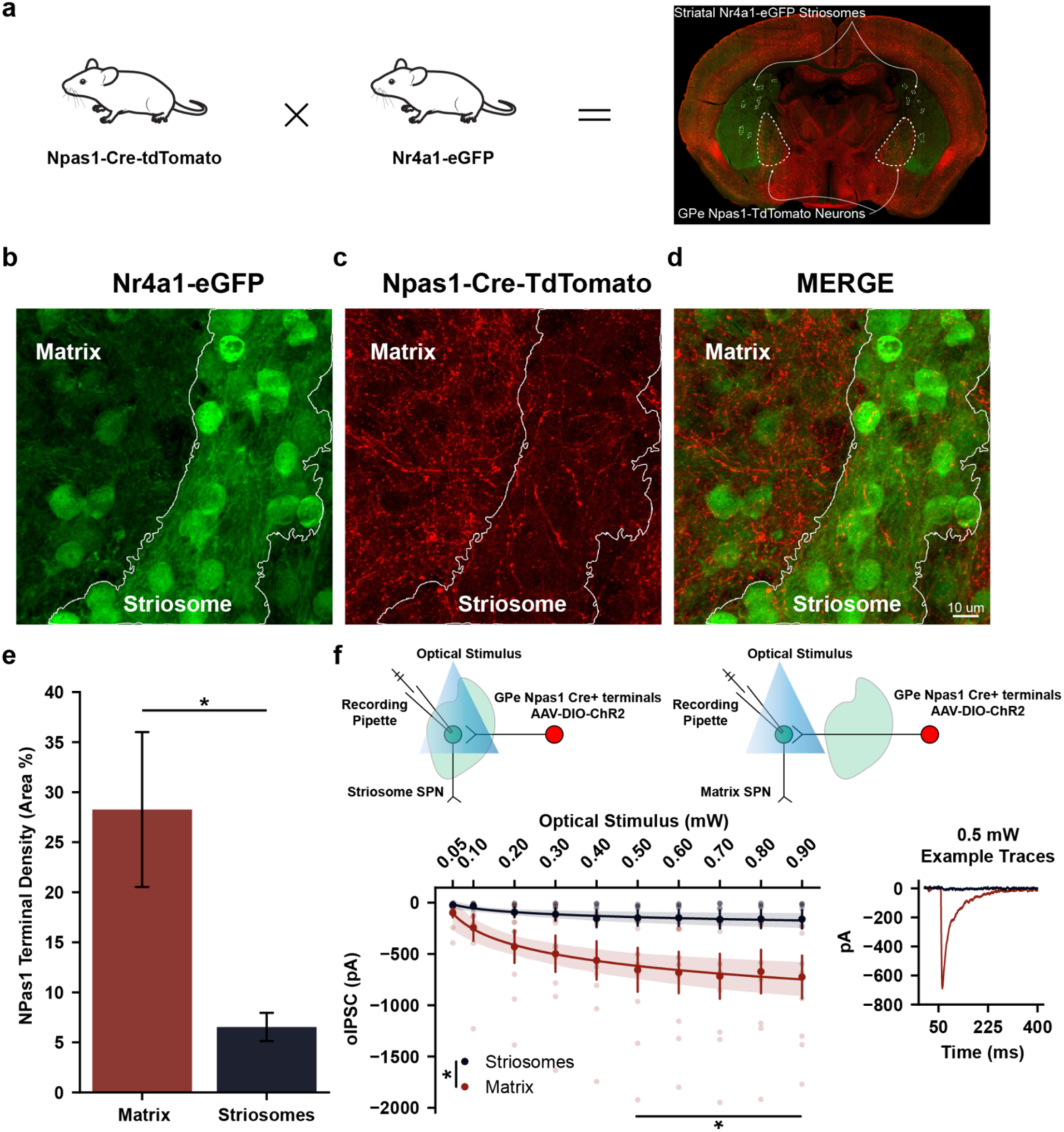
GPe^NPAS1^ neurons preferentially innervate the striatal matrix and exert stronger functional inhibition onto striatal matrix SPNs. **(a)** Representative breeding scheme (Npas1-Cre-tdTomato × Nr4a1-eGFP) and corresponding coronal section of the dorsal striatum and GPe displaying the genetic strategy used to visualize GPe^NPAS1^ projections in striatal matrix and striosome subcompartments. **(b)** Example matrix (sparse eGFP) and striosome (dense eGFP) territories as defined by Nr4a1-eGFP signal. **(c)** Dense NPAS1^+^ (tdTomato) projection terminals in matrix regions and sparse labeling in striosome compartments overlaid onto matrix and striosome annotations. **(d)** Merge of NPAS1^+^ (tdTomato) labeling with matrix/striosome (eGFP) labeling. **(e)** Quantification of NPAS1⁺ terminal density across compartments, expressed as proportion of labeled area within each region of interest (ROI) (matrix/striosome), reveals a significant enrichment of GPe^NPAS1^ projections in matrix compared to striosomes (paired t-test: *t* = -3.231, *p* = 0.04819, *β* = 0.7372). **(f)** Two diagrams displaying how striosome and matrix whole-cell inhibitory postsynaptic currents evoked by optical stimulation (oIPSCs) of Gpe^NPAS1^ terminals were recorded. oIPSCs recorded from matrix and striosome SPNs in response to increasing laser power reveal greater inhibitory drive onto matrix SPNs with an example trace evoked at 0.5 mW demonstrating larger responses in matrix neurons (two-way mixed ANOVA: optical stimulus (mW) × subcompartment: *F*_(9,144)_ = 4.0603, *p* = 0.00012; Matrix: 9 recordings, Striosome: 9 recordings). Dots represent individual data points, error bars or shaded bands represent standard error of the mean (SEM).

To test whether this anatomical preference corresponds to functional differences in inhibitory drive, we recorded optogenetically evoked inhibitory postsynaptic currents (oIPSCs) from principal matrix and striosome SPNs while photoactivating GPe^NPAS1^ terminals in the dorsal striatum. Increasing optical stimulation power produced substantially larger oIPSCs in matrix SPNs compared to striosome SPNs, with clear divergence at stimulation intensities above 0.5 mW (Fig. 1f).

Together, these findings demonstrate that GPe^NPAS1^ neurons selectively innervate and preferentially inhibit matrix SPNs, positioning them to regulate sensorimotor and associative domains of dorsal striatal functions while largely sparing striosomal compartments canonically associated with reward valuation^26^.

### GPe^NPAS1^ manipulations do not alter global performance or locomotion in the elevated plus maze

Given the anatomical and functional specificity of GPe^NPAS1^ terminals within the striatal matrix, we next asked whether modulating these neurons influences behaviors that rely on sensorimotor and associative processing. Although the GPe has traditionally been viewed as a motor output structure, accumulating evidence suggests that its diverse neuronal populations may also contribute to evaluating action-selection behaviors and shaping decision-making strategies^28–31^. To test this possibility in an ethologically relevant context, we used the elevated plus maze (EPM), a task in which animals must physically navigate the environment while continuously assessing the relative risk of its different compartments. Because the EPM intrinsically couples locomotion with threat evaluation, it provides a useful framework for probing motor components and associative decision variables both jointly and, to a degree, separately^32–36^.

To capture these behavioral components with sufficient resolution, we analyzed EPM video recordings using SqueakPose Studio, a deep-learning based pose estimation pipeline that reconstructs full-body posture, heading direction, and locomotor kinematics from freely moving animals^37^. This approach enabled quantification of both coarse measures (*e.g.*, arm occupancy and distance traveled) and fine-grained pose dynamics relevant to risk evaluation. To assess how GPe^NPAS1^ neurons influence these behaviors, we chemogenetically modulated this population by bilaterally expressing Cre-dependent inhibitory hM4D(Gi) or excitatory hM3D(Gq) DREADDs selectively in Npas1⁺ neuron cell bodies within the GPe. Administration of the designer ligand, C21 (1.0 mg/kg), 15 min prior to EPM session which allowed bidirectional chemogenetic manipulation of GPe^NPAS1^ activity while leaving other GPe neuronal subpopulations unperturbed (Fig. 2a).

**Figure 2.**
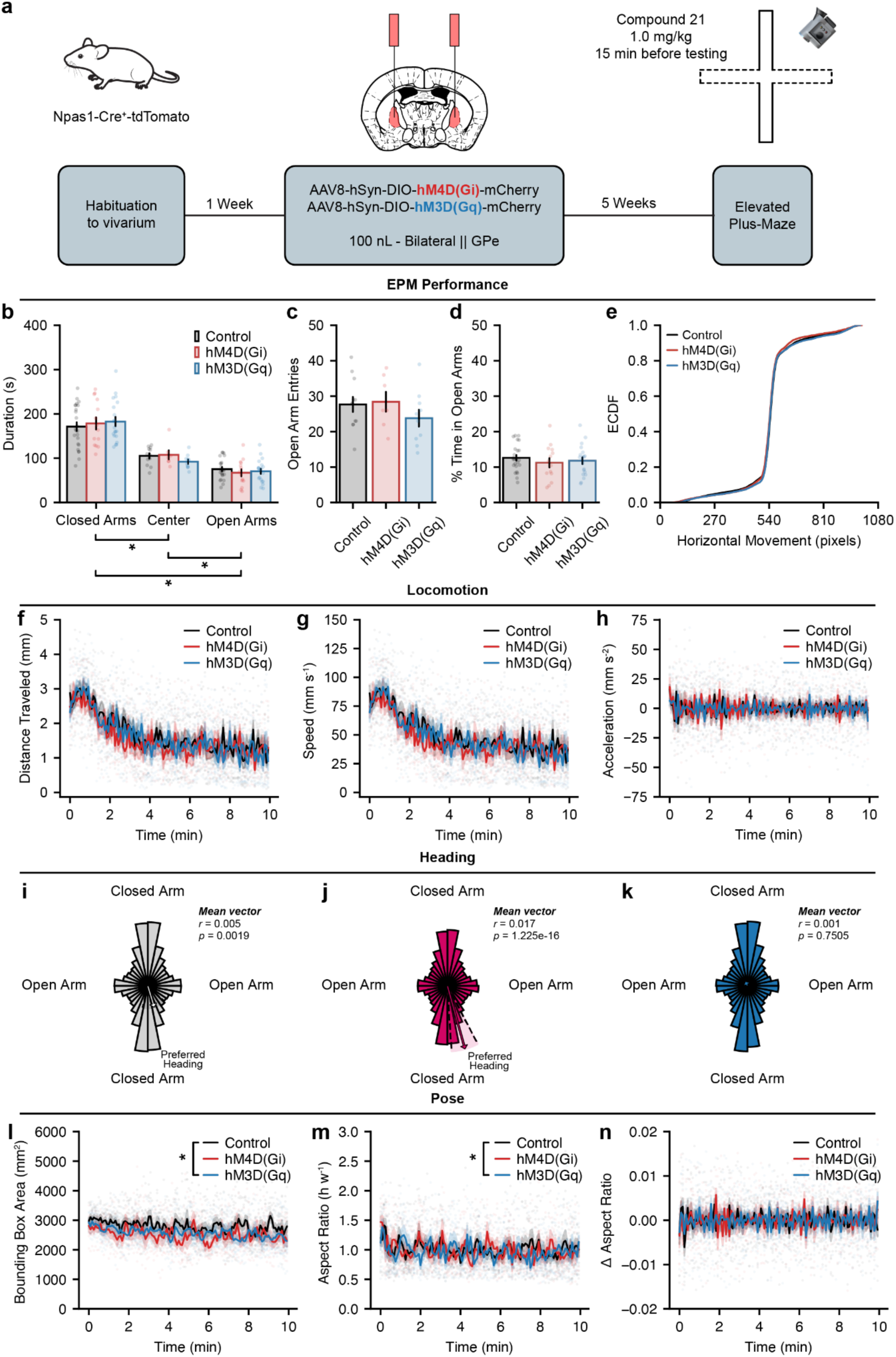
GPe^NPAS1^ manipulations do not alter global performance or locomotion in the elevated plus maze. **(a)** Experimental diagram. Npas1-Cre-TdTomato mice received bilateral GPe injections of Cre-dependent AAV8-hSyn-DIO-hM4D(Gi)-mCherry or AAV8-hSyn-DIO-hM3D(Gq)-mCherry, with Cre-negative littermates serving as controls. All animals received C21 prior to testing, ensuring equivalent drug exposure across groups. Mice were tested on the elevated plus maze (EPM) 5 weeks post-surgery. Behavioral sessions were video-recorded for subsequent analysis. **(b)** Time spent in closed arms, center, and open arms during the EPM shows the expected preference for closed arms across all groups, with no effect of GPe^NPAS1^ manipulation on arm occupancy (two-way mixed ANOVA, maze compartment *F*(2,52) = 122.8, *p* = 2.003e-20; group *F*(2,26) = 1.546, *p* = 0.2319). **(c)** Number of open-arm entries does not differ across groups, indicating no effect of GPe^NPAS1^ manipulation on exploration of these areas (ANOVA, group *F*(2,26) = 1.048, *p* = 0.365). **(d)** Percent time spent in the open arms in the EPM is comparable across control, hM4D(Gi), and hM3D(Gq) mice, consistent with preserved global EPM performance (ANOVA, group *F*(2,55) = 0.4928, *p* = 0.6136). **(e)** Empirical cumulative distribution functions (ECDFs) of frame-wise horizontal movement in the open arms during the EPM show highly overlapping movement distributions across groups, illustrating the absence of gross shifts in locomotor behavior within high-risk regions of the maze. Pairwise Kolmogorov–Smirnov tests detected statistically significant but very small distributional differences (KS D = 0.02–0.05; FDR-corrected *p* < 0.001), consistent with negligible effect sizes that do not reflect meaningful differences in open-arm movement dynamics. **(f)** Distance traveled (two-way mixed ANOVA, group x time *F*(238,3094) = 0.8741, *p* = 0.9131), **(g)** speed (two-way mixed ANOVA, group x time *F*(238,3094) = 0.8742, *p* = 0.9129), and **(h)** acceleration (two-way mixed ANOVA, group x time *F*(238,3094) = 1.037, *p* = 0.3419) over time are comparable across control, hM4D(Gi), and hM3D(Gq) mice, indicating preserved global locomotor output across the session. Frame-wise polar histograms of heading direction during EPM show **(i)** control mice exhibit a modest but significant preference for a closed-arm-oriented heading (Rayleigh test, *r* = 0.005399, *p* = 0.001871). **(j)** hM4D(Gi) mice show a statistically significant, strong preferred closed arm heading (Rayleigh test, *r* = 0.01707, *p* = 1.225e-16). **(k)** In contrast, GPe^NPAS1^ hM4D(Gq) mice do not exhibit a statistically significant preferred heading orientation (Rayleigh test, *r* = 0.001264, *p* = 0.7505). **(l)** Pose features extracted from video tracking show bound box area across time was decreased for hM3D(Gq) mice compared to control mice across all EPM areas (two-way mixed ANOVA, group x time *F*(238,3094) = 1.180, *p* = 0.03497; post hoc control v hM3D(Gq) *p* = 0.03348). **(m)** Similarly, box aspect ratio over time was decreased for hM3D(Gq) mice compared to control mice (two-way mixed ANOVA, group x time *F*(238,3094) = 1.181, *p* = 0.03486; post hoc control v hM3D(Gq) *p* = 0.04464). **(n)** There were no group differences in the change in aspect ratio across time (two-way mixed ANOVA, group x time *F*(238,3094) = 0.08639, *p* = 0.06231). Dots represent individual data points, error bars or shaded bands represent standard error of the mean (SEM). For polar plots, 32 bins were computed to generate 11.25 degree bars for histogram densities.

Across all groups, mice exhibited the expected preference for the closed arms of the EPM, spending significantly more time in closed arms than in the center or open arms, respectively (Fig. 2b). Chemogenetic modulation of GPe^NPAS1^ neurons did not alter open-arm entries or the percentage of time spent in the open arms (Fig. 2c–d). To assess whether locomotor output within high-risk regions of the maze (center and open arms) was altered, we examined frame-wise horizontal movement using empirical cumulative distribution functions (ECDFs). Although statistically detectable differences were observed between groups, the Kolmogorov-Smirnov statistic was extremely small, indicating that overall locomotor output and its distribution were largely preserved and that these differences likely reflect the large number of frame-wise samples and high sensitivity of the K-S statistic rather than biologically meaningful changes in behavior (Fig. 2e). Thus, while gross locomotor behavior in the open arms and center of the EPM was comparable across groups, these analyses do not exclude the possibility that GPe^NPAS1^ manipulation influences the temporal organization or sequencing of exploratory actions within high-risk regions.

To further determine whether GPe^NPAS1^ manipulation affected the temporal dynamics of movement during the task, we examined distance traveled, movement speed, and acceleration as a function of time across the EPM session. All three measures exhibited similar temporal profiles across control, hM4D(Gi), and hM4D(Gq) mice, with no significant group × time interactions observed for any locomotor metric (Fig. 2f–h). These results indicate that neither the magnitude nor the time course of locomotor output - including exploration onset, habituation, or sustained movement - was altered by chemogenetic modulation of GPe^NPAS1^ neurons. Notably, these preserved locomotor dynamics were observed despite the traditional association of the GPe with motor control, suggesting that GPe^NPAS1^ neurons do not contribute strongly to gross movement execution in this context.

To examine whether GPe^NPAS1^ modulation influenced the organization of exploratory orientation independent of global locomotor output, we next analyzed frame-wise heading direction during EPM exploration. Frame-wise heading orientations (a directional vector for where the mouse head was pointing) were plotted with polar histograms and revealed a modest but significant bias in heading orientation in control mice, with headings preferentially oriented toward a closed arm of the maze (Fig. 2i). Inhibition of GPe^NPAS1^ neurons with hM4D(Gi) was associated with a stronger and more consistent closed-arm–oriented heading bias (Fig. 2j), whereas activation of this population with hM4D(Gq) abolished this directional preference, resulting in a more uniform distribution of heading angles (Fig. 2k). Together, these results indicate that although global locomotor dynamics are preserved, modulation of GPe^NPAS1^ neurons alters the structure of orienting behavior.

Finally, to further increase the resolution with which we assessed global behavior during EPM exploration, we analyzed pose features extracted from frame-wise video tracking. Bounding box area (the area of a rectangular box drawn around the perimeter of the mouse, excluding the tail) exhibited a modest group-dependent effect across the session, with a significant group × time interaction and reduced bounding box area in hM4D(Gq) mice relative to controls - demonstrating that Gq treated mice preferentially spent more time in constrained, non-stretched body postures (Fig. 2l). Body aspect ratio (as measured by the height of the bounding box divided by the width) similarly differed in hM4D(Gq) mice relative to controls, indicating a stable shift in postural configuration associated with GPe^NPAS1^ modulation (Fig. 2m). In contrast, moment-to-moment (frame-wise differences from one frame to the next) changes in aspect ratio were preserved across groups, with no significant group or group × time effects detected (Fig. 2n). Importantly, the absence of main effect group differences in postural dynamics, together with preserved locomotor output across time, argues against trivial explanations such as differences in body size or movement vigor that explain the observed postural differences. Instead these data suggest that GPe^NPAS1^ manipulation influences the postural configurations animals adopt during exploration at certain moments in time.

Together, these analyses demonstrate that chemogenetic modulation of GPe^NPAS1^ neurons does not alter canonical measures of EPM performance or gross locomotor output, including arm occupancy, open-arm exploration, movement magnitude, or the temporal dynamics of locomotion. Despite preserved global behavior, higher-resolution analyses revealed subtle but structured differences in orientation and postural configuration, including altered heading biases and divergence in bounding box area and aspect ratio at specific moments in time. These findings indicate that GPe^NPAS1^ manipulation does not influence whether animals explore risky environments, but rather how exploratory actions may be organized in space and time. We therefore next examined whether GPe^NPAS1^ neurons regulate the sequencing and resolution of risk-related exploratory behaviors during EPM exploration.

### GPe^NPAS1^ neurons shape the structure and sequencing of risk-ranked behaviors

To determine whether the orientation and postural differences observed in Figure 2 reflect altered organization of exploratory actions, we analyzed the structure and sequencing of risk-related behaviors during EPM exploration. Rather than treating EPM performance as a collection of independent spatial metrics (*e.g.*, closed v open arm occupancy), we segmented the continuous 10-minute EPM session into discrete behavioral sequences defined by the order and resolution of arm-to-arm transitions (Fig. 3a). This approach allowed us to quantify not only where animals traveled, but how they progressed through each behavioral sequence under varying degrees of risk.

**Figure 3.**
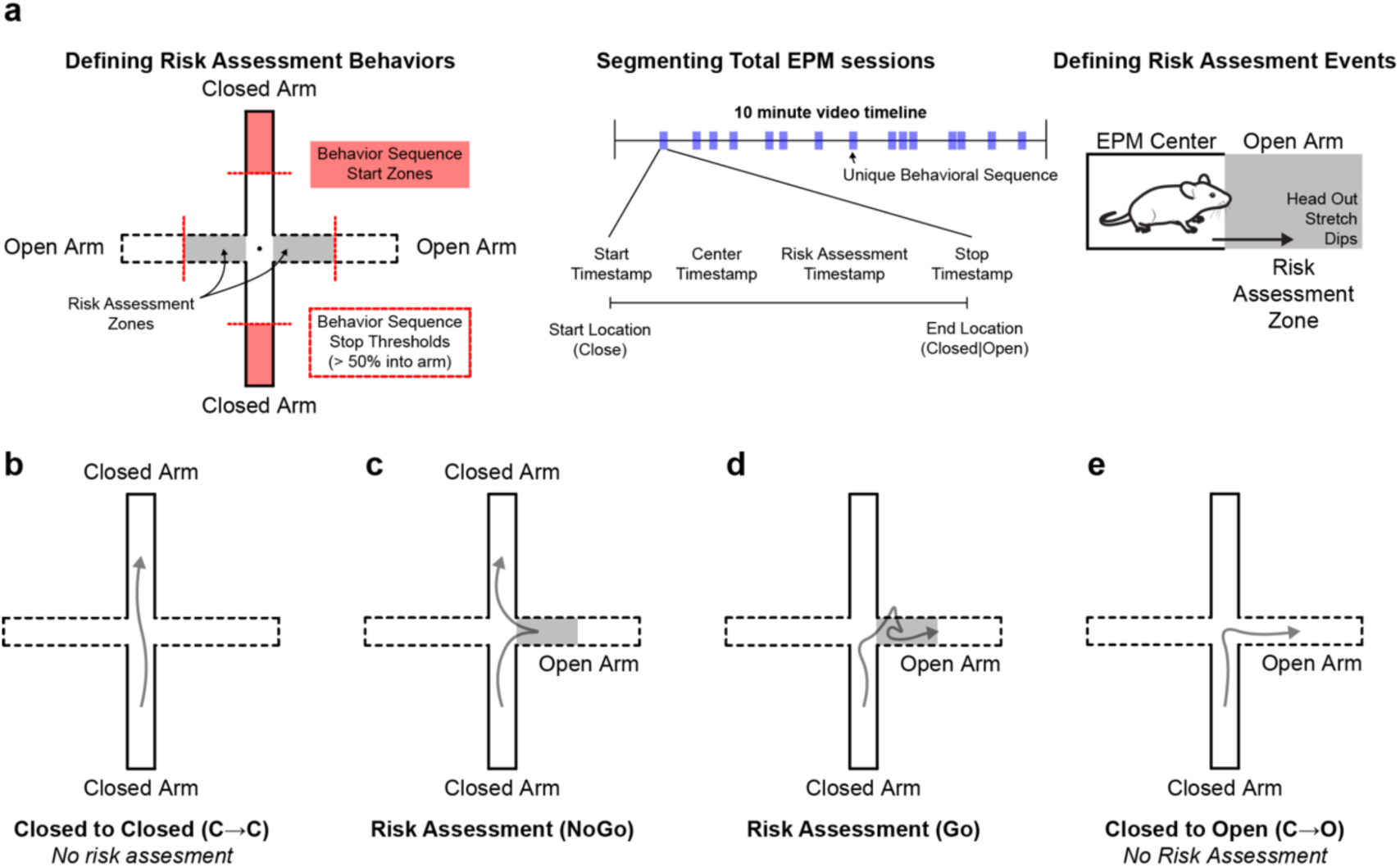
Segmenting EPM behaviors into discrete risk-ranked behavioral sequences. **(a)** Diagram illustrating the rule-based segmentation of continuous elevated plus maze (EPM) behavior into discrete risk-ranked behavioral sequences. Sequences were initiated when animals occupied the distal portion of a closed arm (sequence start zones) and terminated upon crossing ≥50% into a subsequent arm (sequence stop threshold). Risk assessment behaviors were defined by stereotyped head-out postures, body stretches, or edge dips in the proximal region of the open arms before crossing the sequence stop threshold. Sequences were manually annotated using BORIS to generate start, transition (center), risk assessment, and stop timestamps. **(b)** Closed to Closed (C→C) sequences occurred when mice started and ended in the closed arms of the EPM without sampling the open arm. **(c)** NoGo sequences occurred when mice started in the closed arm, performed at least 1 risk assessment behavior in the proximal region of the open arm, and then returned to a closed arm. **(d)** Go sequences occurred when mice started in the closed arm, performed at least 1 risk assessment behavior in the proximal region of the open arm, and then crossed the threshold to end the sequence on the distal end of the open arm. **(e)** Closed to open (C→O) sequences occurred when animals started in the closed arm, traversed the proximal part of the open arm without performing any risk assessment behaviors, and ended the sequence in the distal part of the open arm.

To define these discrete risk-ranked sequences, we initiated sequence starts when mice were in the distal part of the closed arm (start sequence zones) with headings and velocities oriented towards the center of the EPM. When animals crossed beyond 50% into the non-originating arm (stop thresholds) we labeled the completion of the risk-ranked sequence. We identified four distinct sequence types: direct closed-to-closed transitions (C→C) (Fig. 3b), risk assessment sequences beginning and terminating in the closed arms (NoGo) (Fig. 3c), risk assessment sequences beginning in closed arms and culminating in open-arm entry (Go) (Fig. 3d), and direct closed-to-open transitions (C→O) that occurred without any risk assessment behavior in the proximal part of the open arm (Fig. 3e). These sequence types reflect increasing commitment to risk, respectively and dissociate different types of action execution. For example behavior videos of each discrete risk-ranked behavior, please see supplementary data files 1 - 4.

We first examined closed-to-closed (C→C) sequences, which represent ongoing exploration confined to the closed arms without engagement of the open arms. Trajectory maps confirmed that these sequences were spatially restricted to the closed arms and exhibited stereotyped movement patterns across all groups that represent the least amount of exploratory risk (Fig. 4a). Quantification of sequence features revealed group-dependent differences in temporal structure: activation of GPe^NPAS1^ neurons (hM4D(Gq)) was associated with shorter C→C sequence durations compared to control and hM4D(Gi) mice, whereas no difference was observed between control and hM4D(Gi) groups (Fig. 4b). These effects were accompanied by corresponding changes in average movement speed, while average aspect ratio within C→C sequences remained preserved across groups (Fig. 4c–d), indicating that GPe^NPAS1^ modulation primarily influenced the timing and kinematics of closed-arm exploration rather than its postural expression.

**Figure 4.**
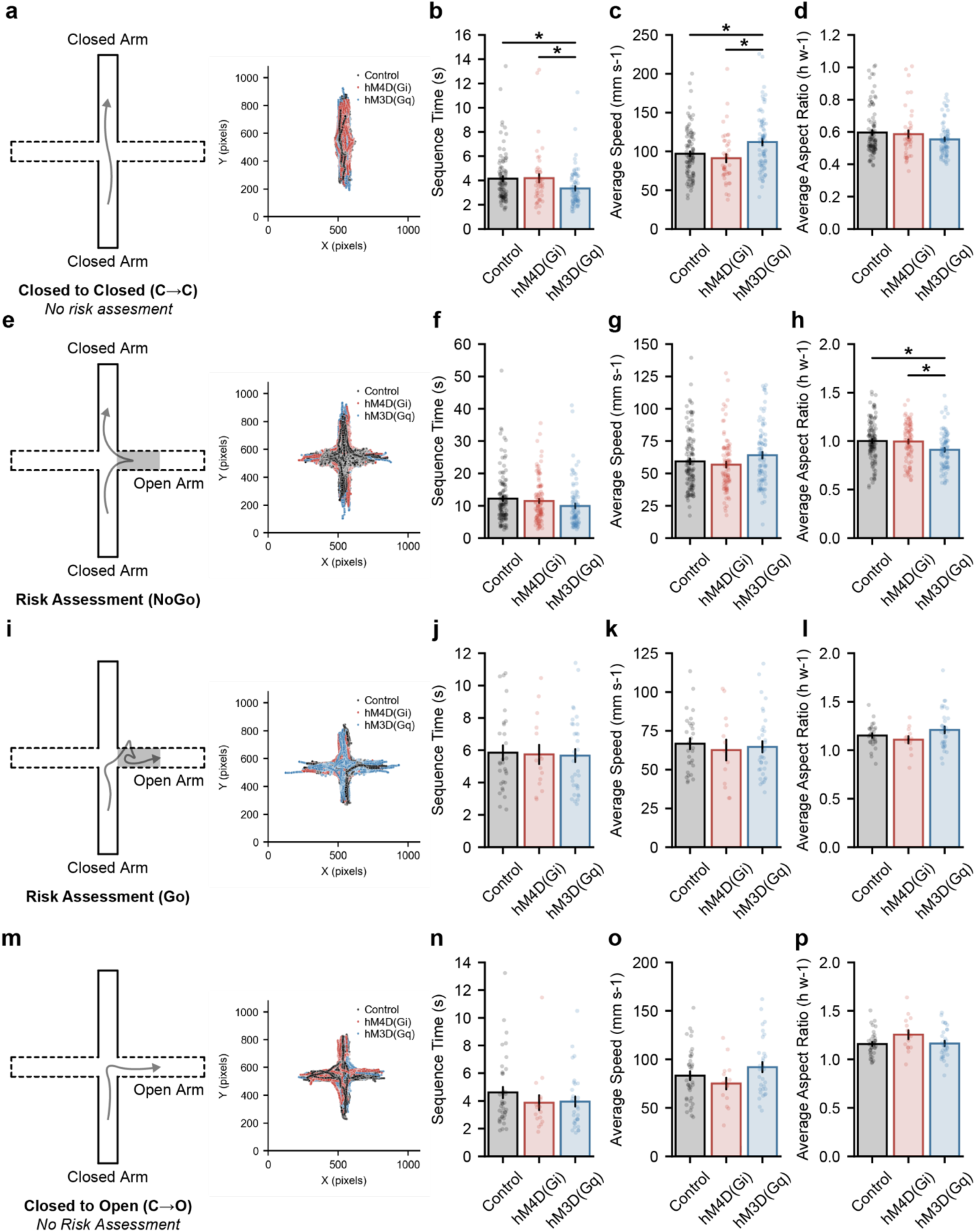
GPe^NPAS1^ modulation alters the structure of risk-ranked behavior sequences. **(a)** Trajectory maps showing frame-wise tracking data for closed to closed (C→C) sequences, illustrating exploration confined to the closed arms across all groups. **(b)** Activation of GPe^NPAS1^ neurons (hM4D(Gq)) was associated with shorter C→C sequence durations compared to control and hM4D(Gi) mice (ANOVA, group *F*(2,216) = 4.368, *p* = 0.01381; post hoc control v Gq *p* = 0.003385, Gi v Gq *p* = 0.03239). **(c)** Activation of GPe^NPAS1^ neurons (hM4D(Gq)) was associated with faster average C→C sequence speed compared to control and hM4D(Gi) mice (ANOVA, group *F*(2,211) = 6.473, *p* = 0.001869; post hoc control v Gq *p* = 0.005505, Gi v Gq *p* = 0.002394). **(d)** No group differences were observed in average aspect ratio during C→C sequences (ANOVA, group *F*(2,211) = 2.170, *p* =0.1166). **(e)** Trajectory maps showing frame-wise tracking data for risk assessment (NoGo) sequences, illustrating exploration in risk assessment regions without full open-arm entry. **(f)** No group differences were observed in sequence time (ANOVA, group *F*(2,314) = 2.557, *p* = 0.07907) and **(g)** average speed (ANOVA, group *F*(2,297) = 2.438, *p* = 0.08904) during NoGo sequences. **(h)** Activation of GPe^NPAS1^ neurons (hM4D(Gq)) was associated with smaller aspect ratios during NoGo sequences compared to control and hM4D(Gi) mice (ANOVA, group *F*(2,297) = 6.322, *p* = 0.002047; post hoc control v Gq *p* = 0.00098, Gi v Gq *p* = 0.003128). **(i)** Trajectory maps showing frame-wise tracking data for risk assessment (Go) sequences, illustrating exploration in risk assessment regions with full open-arm entry. No group differences were observed for **(j)** Go sequence duration (ANOVA, group *F*(2,75) = 0.04263, *p* = 0.9582), **(k)** Go sequence speed (ANOVA, group *F*(2,73) = 0.1923, *p* = 0.8254), **(l)** or Go sequence posture (ANOVA, group *F*(2,73) = 2.076, *p* = 0.1327). **(m)** Trajectory maps showing frame-wise tracking data for closed to open (C→O) sequences, illustrating direct closed to open transitions absent risk assessment. No group differences were observed for **(n)** C→O sequence duration (ANOVA, group *F*(2,84) = 0.9651, *p* = 0.3851), **(o)** C→O sequence speed (ANOVA, group *F*(2,82) = 2.051, *p* = 0.8254), **(p)** or C→O sequence posture (ANOVA, group *F*(2,82) = 2.451, *p* = 0.09244). Dots represent individual data points, error bars represent standard error of the mean (SEM).

We next examined risk assessment (NoGo) sequences, defined as exploration beginning in the closed arms, followed by engagement of risk-assessment behaviors (*e.g.*, head-out postures, body stretches, and edge dips) in the proximal region of the open arm and subsequent withdrawal back into the closed arms without crossing the ≥ 50% open-arm threshold. Trajectory maps confirmed that NoGo sequences involved excursions into the proximal regions of the open arms without full open-arm entry (Fig. 4e). Average sequence duration and movement speed during NoGo events were comparable across control, hM4D(Gi), and hM4D(Gq) mice (Fig. 4f–g). Notably, NoGo sequences were longer than closed-to-closed transitions across all groups, consistent with the extended evaluation period associated with risk assessment. Despite preserved timing and kinematics, activation of GPe^NPAS1^ neurons (hM4D(Gq)) was associated with smaller aspect ratios during NoGo sequences compared to control and hM4D(Gi) mice (Fig. 4h), indicating altered postural configuration during risk assessment behaviors.

In contrast, higher-risk sequences, including risk assessment sequences that culminated in open-arm entry (Go) and direct closed-to-open transitions (C→O), showed no group-dependent differences. Trajectory maps confirmed stereotyped approaches and full open-arm entry across all groups (Fig. 4i,m). Sequence duration, movement speed, and postural configuration during Go and C→O sequences were overlapping across control, hM4D(Gi), and hM4D(Gq) mice (Fig. 4j–l, n–p), suggesting that once animals committed to open-arm exploration, execution of these high-risk actions was relatively preserved for the variables measured.

To examine how GPe^NPAS1^ modulation influences overall behavioral strategy, we next quantified the distribution of sequence types across the full 10-minute EPM session. The number of all risk-ranked sequences were statistically different from one another, except Go and C→O. And for NoGo sequences, Gi mice had an increased number of NoGo sequences compared to control and Gq mice (Fig. 5a). These data further suggest that lower risk behaviors (e.g. C→C and NoGo) are distinct from higher risk behaviors and that modulating GPe^NPAS1^ neurons can drive measurable differences in action-selection behaviors. Because these behaviors also occur along a continuum of risk engagement, we computed a risk assessment index defined as the proportion of sequences involving risk assessment (NoGo + Go) relative to total sequences. Although hM4D(Gi) mice did not exhibit changes in sequence timing, kinematics, or posture compared to controls (Fig. 4b-d) - they displayed a higher proportion of risk assessment sequences compared to control and hM4D(Gq) mice (Fig. 5b). In contrast, hM4D(Gq) mice, despite showing altered timing and postural configuration during low-risk (*e.g.*, C→C and NoGo) sequences, exhibited a lower proportion of risk assessment behaviors relative to hM4D(Gi) mice, but not controls (Fig. 5b). Together, these results indicate that inhibition of GPe^NPAS1^ neurons selectively increases the likelihood of engaging in risk assessment behaviors without altering their execution, whereas activation of this population alters the timing and postural configuration of exploratory sequences, particularly at lower levels of risk (NoGo).

**Figure 5.**
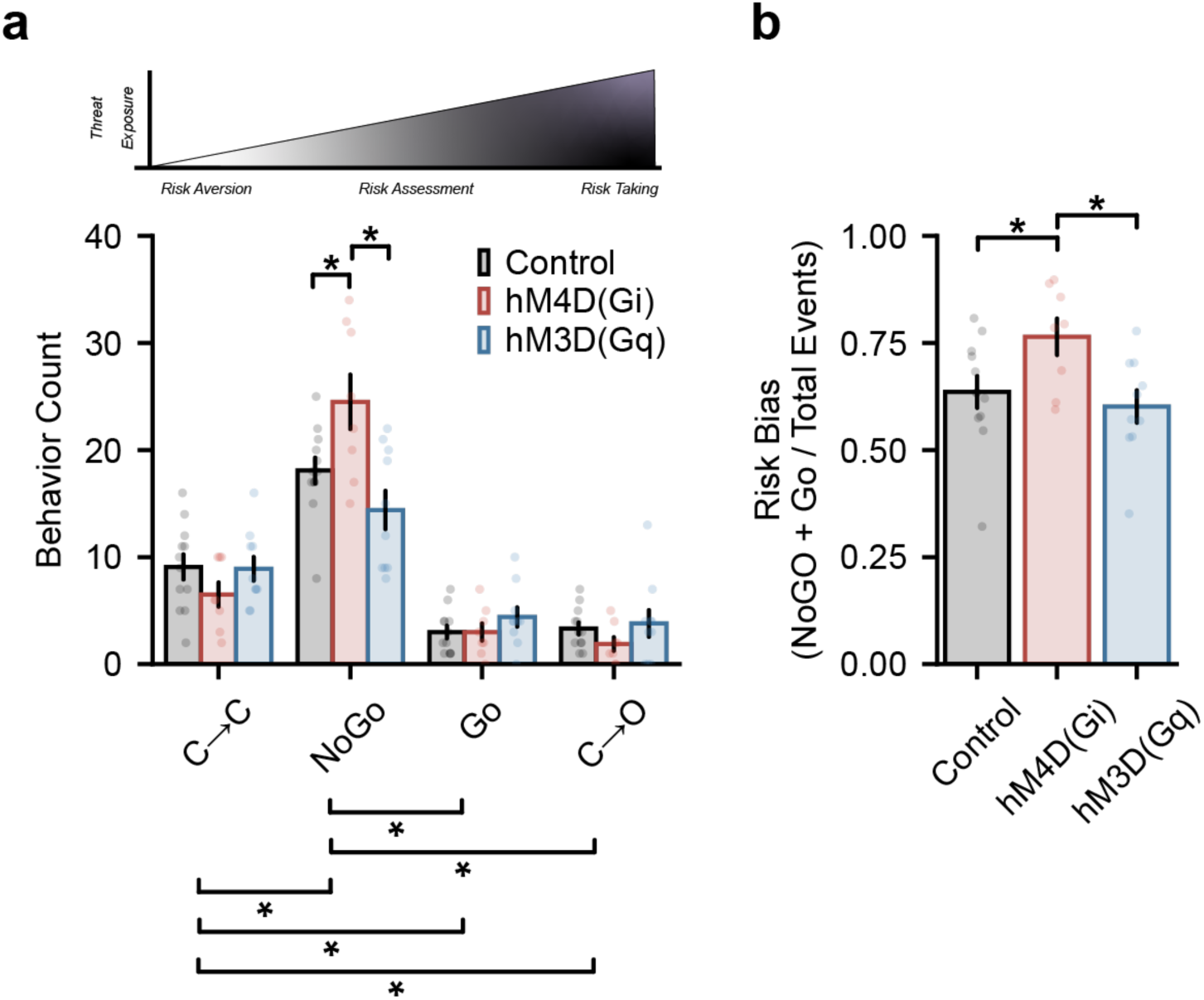
GPe^NPAS1^ modulation biases the frequency of risk assessment sequence selection. **(a)** Sequence counts by type across the 10-minute elevated plus maze (EPM) session show shifts in the frequency of NoGo behaviors for hM4D(Gi) compared to controls and hM3D(Gq). (two-way mixed ANOVA, group x sequence type *F*(6,81) = 5.334, *p* = 0.00011; post hoc NoGo: Control v Gi *p* = 0.0453, NoGo: Gq v Gi *p* = 0.0063) **(b)** Risk assessment index, calculated as the proportion of sequences involving risk assessment behaviors (NoGo + Go) relative to the total number of sequences observed per animal. hM4D(Gi) mice exhibited an increased proportion of risk assessment sequences compared to controls, whereas hM4D(Gq) mice showed a reduced proportion of risk assessment sequences relative to hM4D(Gi) mice, with values comparable to controls (ANOVA, group *F*(2,27) = 4.193, *p* = 0.02594; post hoc control v Gi *p* = 0.03691, Gi v Gq *p* = 0.01202, Gq v control *p* = 0.5289). Dots represent individual data points, error bars represent standard error of the mean (SEM).

### Presynaptic activity of NPAS1 neurons show sequence- and outcome-specific activity dynamics during risk transitions

Chemogenetic manipulation of GPe^NPAS1^ neurons perturbs activity of neuronal somata and all downstream projections and exerts continuous effects throughout the EPM session. Thus, the behavioral changes observed in Figures 2–5 do not directly reveal how arkypallidal GPe^NPAS1^ presynaptic activity is engaged during specific risk-related exploratory sequences. To more directly map sequence-level behavioral changes onto neural activity, we expressed the calcium indicator GCaMP8f selectively in GPe^NPAS1^ neuron cell bodies and implanted fiber optic probes bilaterally in the dorsal striatum, enabling recording of arkypallidal GPe^NPAS1^ presynaptic terminal activity during EPM exploration (Fig. 6a).

**Figure 6.**
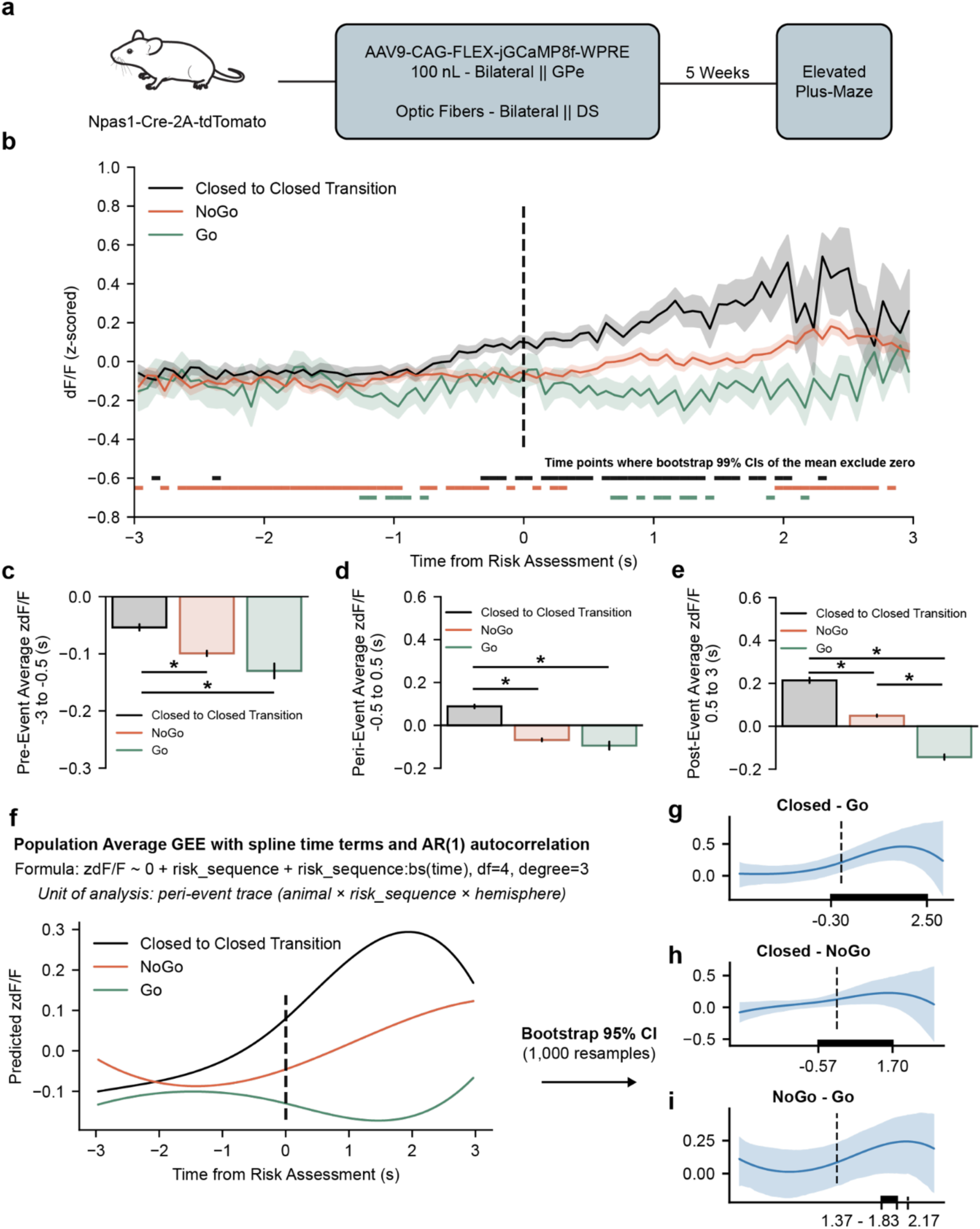
Arkypallidal NPAS1 terminals in the striatal matrix show sequence- and outcome-specific activity dynamics during risk transitions. **(a)** Npas1-Cre mice received bilateral GPe injections of AAV9-CAG-FLEX-jGCaMP8f-WPRE and bilateral optical fiber implants in dorsal striatum (DS). Five weeks after surgery, mice were tested on the elevated plus maze (EPM) while GPe^NPAS1 terminal activity was recorded. **(b)** Event-aligned GPe^NPAS1^ terminal calcium activity (z-scored ΔF/F) during closed-to-closed transitions (C→C), risk assessment that did not culminate in open-arm entry (NoGo), and risk assessment culminating in open-arm entry (Go). NoGo and Go traces were aligned to the onset of the first risk assessment; C→C traces were aligned to center crossing (time 0, dashed line). Traces show the mean across peri-event units (animal × sequence type × hemisphere), with shaded regions indicating ±SEM across units. Colored horizontal bars indicate time points where bootstrap 99% confidence intervals of the mean excluded zero for the corresponding sequence type. **(c)** Mean pre-event GPe^NPAS1^ terminal activity (−3 to −0.5 s relative to event onset) shows decreases in NoGo and Go compared to C→C sequences (ANOVA, behavior sequence *F*(2,25469) = 26.37, *p* = 3.598e-12; post hoc C→C v Go *p* = 2.034e-7, C→C v NoGo *p* = 1.064e-9, NoGo v Go *p* = 0.07621). **(d)** Mean peri-event GPe^NPAS1^ terminal activity (−0.5 to 0.5 s relative to event onset) shows decreases in NoGo and Go compared to C→C, with C→C zdF/F values moving from negative in the pre-event window to positive in the peri-event window (ANOVA, behavior sequence *F*(2,11945) = 123.1, *p* = 1.139e-53; post hoc C→C v Go *p* = 6.733e-19, C→C v NoGo *p* = 1.559e-49, NoGo v Go *p* = 0.4397). **(e)** Mean post-event GPe^NPAS1^ terminal activity (0.5 to 3 s relative to event onset) shows difference between all sequence types, with C→C the highest, positive GPe^NPAS1^ terminal calcium activity, NoGo with slightly positive activity, and Go with negative activity (ANOVA, behavior sequence *F*(2,20093) = 191.8, *p* = 3.061-83; post hoc C→C v Go *p* = 1.678e-88, C→C v NoGo *p* = 4.136e-34, NoGo v Go *p* = 8.636e-46). (f) *Population-average modeling.* Sequence-specific time courses were modeled using generalized estimating equations (GEE) with cubic B-spline time terms (df = 4) and AR(1) autocorrelation, fit at the level of individual peri-event units (animal × sequence type × hemisphere; n = 612 units from 14 mice). Model-predicted z-scored dF/F traces are shown (1,000 resamples). The fitted AR(1) correlation parameter was ρ = 0.987. A joint Wald test of all spline interaction terms indicated a significant time × sequence interaction (Wald χ²(15) = 102.9, p = 3.69e−15), confirming that zdF/F time courses differed across sequence types. Pairwise differences in modeled time courses were shown by subtracting mouse-level bootstrap 95% confidence bands across the event window to determine where in time behavioral sequence signals differed. **(g)** Increases in C→C GPe^NPAS1^ terminal activity were seen relative to Go sequences from -3 to 2.5 s (Wald χ²(4) = 28.07, *p* = 1.209e-5). **(h)** Increases in C→C GPe^NPAS1^ terminal activity were also seen relative to NoGo sequences, but during a left-shifted, shorter duration from -0.57 to 1.7 s (Wald χ²(4) = 24.63, *p* = 5.957e-5). **(i)** Small increases in NoGo GPe^NPAS1^ terminal activity relative to Go sequences were observed in the post event window from 1.37 to 1.83 s and at 2.17 s (Wald χ²(4) = 14.37, *p* = 0.006187).

Using the same rule-based behavioral segmentation described in Figure 3, we timestamped sequence initiation, center crossings, risk assessment onset, and sequence termination for each behavioral event. For risk assessment sequences (Go and NoGo), photometry signals were aligned to the onset of the first risk assessment behavior, whereas C→C transitions were aligned to the time at which animals crossed the center of the maze (Fig. 6b). Event-aligned photometry traces were normalized within animal and then analyzed across a six-second peri-event window.

Event-aligned recordings revealed clear sequence-specific dynamics in GPe^NPAS1^ terminal activity in the dorsal striatum. C→C transitions were associated with a progressive increase in terminal activity preceding and following the transition, whereas NoGo sequences exhibited a smaller post-event increase. In contrast, Go sequences were characterized by sustained suppression of GPe^NPAS1^ terminal activity throughout the risk assessment and commitment period (Fig. 6b). Consistent with this pattern, pre-event activity (-3 to -0.5 s) was modestly negative across all sequence types and became increasingly suppressed with greater risk engagement (Fig. 6c). During the peri-event window (−0.5 to 0.5 s), C→C transitions exhibited higher GPe^NPAS1^ terminal activity relative to both NoGo and Go sequences (Fig. 6d). In the post-event window, activity increased for C→C and NoGo sequences, whereas GPe^NPAS1^ terminal activity remained suppressed following Go risk assessment (Fig. 6e).

To quantify these sequence-dependent dynamics while accounting for repeated peri-event traces nested within animals, photometry signals were modeled using a generalized estimating equation (GEE) framework with spline-based time terms and AR(1) autocorrelation. Model-derived estimates revealed sustained elevations in GPe^NPAS1^ terminal activity following C→C transitions relative to Go sequences from approximately −0.3 to 2.5 s after the event (Fig. 6g). C→C transitions also exhibited higher activity than NoGo sequences over a shorter post-event interval (Fig. 6h), whereas NoGo sequences showed modestly elevated activity relative to Go sequences for a brief period following risk assessment onset (Fig. 6i).

Together, these results demonstrate that arkypallidal GPe^NPAS1^ terminal activity in the dorsal striatum encodes the structure and resolution of risk-related exploratory sequences, with activity highest during low-risk state persistence (C→C), intermediate during risk assessment without commitment (NoGo), and suppressed during commitment to open-arm exploration (Go). Importantly, although time-averaged pre-event analyses suggested modest sequence differences, modeling the full event-aligned time course revealed no significant pre-event divergence across sequence types, indicating that sequence-specific differentiation in GPe^NPAS1^ terminal activity emerges primarily during peri- and post-event epochs.

## Discussion

Adaptive exploration requires animals to continuously balance information gathering with threat avoidance, a process that depends not only on evaluating risk but also on flexibly transitioning between behavioral states. By combining anatomical mapping, electrophysiology, chemogenetic manipulation, and *in vivo* calcium imaging during EPM exploration, we show that GPe^NPAS1^ neurons preferentially innervate the matrix subcompartment of the dorsal striatum and, under these conditions, do not regulate gross locomotor output. Instead, they shape how exploratory actions unfold across time and how these actions resolve under uncertainty. Npas1-expressing neurons of the GPe are therefore sufficient to play a key role in regulating the structure, sequencing, and resolution of risk-related exploratory behavior.

A central anatomical finding of this study is that GPe^NPAS1^ neurons preferentially innervate the striatal matrix rather than striosomal subcompartments. This distinct projection pattern positions arkypallidal GPe^NPAS1^ neurons to influence sensorimotor and associative processing domains of the dorsal striatum while largely sparing, relative to matrix, striosome-associated circuits implicated in reward valuation and reinforcement learning. Functionally, this organization suggests that GPe^NPAS1^ neurons are well suited to regulate behavioral sequencing and state transitions rather than encoding value or outcome expectancy *per se*. Consistent with this interpretation, optogenetic activation of GPe^NPAS1^ terminals produced stronger inhibition of matrix spiny projection neurons, providing a synaptic mechanism by which pallidal feedback could modulate the timing and organization of striatal output during ongoing behavior.

Prior work has established that arkypallidal GPe neurons are well positioned to coordinate specialized stop processes in the brain^38–40^. Others have demonstrated that optogenetic excitation of arkypallidal neurons can robustly inhibit ongoing locomotion, and that disruption of the arkypallidal projections can shift habit-seeking behaviors^41,42^. To date, arkypallidal projections contributions to inhibitory control have largely been conceptualized as cancelling an ongoing action^43^. Many of these studies rely on Go/NoGo, Stop Signal, or other cued paradigms in which locomotor performance is tightly coupled to an explicit instruction or trigger signaling the appropriate behavioral response. While these task structures have been invaluable, here we intentionally examined how arkypallidal GPe neurons respond to environmentally driven, task-free changes in action selection, allowing us to dissociate associative and locomotor components of behavior.

Our anatomical and functional connectivity data suggest that GPe^NPAS1^ neurons are better positioned to bias the associative components and competitive dynamics of action-selection, rather than to directly alter locomotor output. By preferentially targeting matrix spiny projection neurons, GPe^NPAS1^ terminals can reshape the race conditions between canonical Go and NoGo pathways that determine how exploratory actions are sequenced, sustained, or terminated. In light of the increasing complexity of basal ganglia circuit organization^44^, these findings argue that pallidal firing should be understood not solely as a modulator of motor execution, but as a regulator of the temporal structure and resolution of behavior.

At the behavioral level, chemogenetic modulation of GPe^NPAS1^ neurons revealed a dissociation between global performance metrics and fine-grained behavioral organization. Neither inhibition nor activation of this population altered canonical EPM measures, including arm occupancy, open-arm entries, or overall locomotor dynamics. Instead, higher-resolution analyses uncovered structured changes in orientation, posture, and the temporal properties of exploratory sequences, particularly during low- and intermediate-risk states. Inhibition of GPe^NPAS1^ neurons increased the likelihood of engaging in risk assessment behaviors without altering their execution, whereas activation of this population reshaped the timing and postural configuration of exploratory actions. These findings indicate that arkypallidal GPe^NPAS1^ neurons bias the selection and organization of behavioral sequences rather than directly suppressing or promoting risky exploration.

Importantly, *in vivo* calcium imaging revealed that GPe^NPAS1^ terminal activity encodes sequence-and outcome-specific dynamics during risk transitions. C→C transitions, which reflect persistence in low-risk states, were associated with rising terminal activity, whereas risk assessment without commitment (NoGo) exhibited intermediate dynamics. In contrast, commitment to open-arm exploration (Go) was marked by sustained suppression of GPe^NPAS1^ terminal activity. Modeling the full event-aligned time course using a GEE framework demonstrated that these differences reflect distinct temporal profiles rather than baseline offsets, with divergence emerging primarily during peri- and post-event epochs. Together, these data suggest that GPe^NPAS1^ activity signals the stability or destabilization of ongoing behavioral states, with suppression accompanying commitment to higher-risk actions.

The relationship between GPe^NPAS1^ terminal activity and risk-related behavior observed *in vivo* provides a conceptual framework for interpreting the chemogenetic manipulations. The observed peri- and post-event suppression of GPe^NPAS1^ terminal activity accompanying higher-risk behavioral sequences is consistent with the observation that inhibition of GPe^NPAS1^ neurons using Gi-coupled DREADDs may bias animals toward higher-risk behavioral states by promoting a low-activity regime that, under natural conditions, is associated with risk assessment and commitment. Rather than inducing risk-taking directly, reducing GPe^NPAS1^ output may shift the system into a dynamic regime that favors risk assessment and commitment, consistent with the increased risk assessment index observed following Gi-mediated inhibition.

In contrast, activation of GPe^NPAS1^ neurons using Gq-coupled DREADDs did not produce a complementary change in risky behavior, despite clear effects on spatial kinematics and posture. One possible explanation is that elevating the baseline activity of this intrinsically autonomical firing neuronal population compresses its dynamic range, impairing the relative activity modulations observed during naturalistic behavior. In this scenario, increasing tonic activity may not enhance the sequence-specific signaling captured by photometry, but instead disrupts the temporal structure of motor output, resulting in altered orientation and posture without promoting risk-related transitions. Together, these findings suggest that GPe^NPAS1^ neurons regulate risk-related behavior through state-dependent modulation of activity dynamics rather than simple increases or decreases in firing rate, highlighting the importance of relative, temporally structured signaling in pallidal control of exploratory behavior.

These results extend emerging views of the GPe as a dynamic regulator of action selection rather than a passive relay in basal ganglia circuitry. While much prior work has emphasized the role of pallidal output in motor control, our findings indicate that arkypallidal GPe^NPAS1^ neurons also contribute to cognitive aspects of behavior by gating transitions between exploratory and defensive states. This function aligns with theoretical frameworks in which basal ganglia circuits regulate not only action initiation but also the timing, sequencing, and termination of behavioral programs under uncertainty. By selectively targeting the striatal matrix, GPe^NPAS1^ neurons may influence how contextual and sensorimotor information is integrated to determine whether animals resist, evaluate, or commit during exploration.

Several limitations and future directions warrant consideration. First, fiber photometry provides population-level signals and cannot resolve heterogeneity within the GPe^NPAS1^ terminal population or distinguish between matrix or striosomal inputs driving observed activity patterns. Future work focused on resolving projection-specific manipulations in the striatum could clarify if there exists functional or timing differences in striosome or matrix targeted arkypallidal GPe^NPAS1^ engagement. Second, while we know that GPe^NPAS1^ somata exhibit spontaneous firing, we know less about the activity of their axons and axon terminals, and thus the changes in activity and effects of modulation may be more complex than we hypothesize. Third, while the EPM provides an ethologically relevant framework for studying risk, it remains to be determined whether similar pallidal dynamics govern decision-making in other contexts involving uncertainty or conflict in a similar task-free environment.

In summary, our findings identify arkypallidal GPe^NPAS1^ neurons as a circuit substrate that shapes the temporal organization of risk-related exploratory behavior. By encoding and modulating behavioral state transitions rather than global action output, this pallidal pathway provides a mechanism through which the basal ganglia can flexibly regulate exploration under uncertainty.

## Methods

### Animals

All experimental procedures were conducted in accordance with institutional ethical guidelines and national regulations and complied with the NIH Guide for the Care and Use of Laboratory Animals (NIH, 2011). Animal experiments were performed at both the Universidade Federal de São Paulo (UNIFESP) and the National Institutes of Health (NIH). Procedures conducted at UNIFESP were approved by the Comissão de Ética no Uso de Animais (CEUA #7826300522; Universidade Federal de São Paulo, Brazil), and procedures conducted at the NIH were reviewed and approved by the NIAAA Animal Care and Use Committee in protocols outlined in animal protocol LIN-DL-1.

Behavioral chemogenetic experiments were performed using Npas1-Cre-2A-tdTomato-BAC mice (12-14 weeks old at arrival; n = 30), a transgenic line expressing Cre recombinase and the tdTomato reporter in Npas1-expressing neurons. For simplification, this line is hereafter referred to as Npas1-Cre-tdTm. Both Cre-positive and Cre-negative littermates were used in the experiments, with Cre-negative mice serving as controls. All animals were obtained from the Centro de Desenvolvimento de Modelos Experimentais para Medicina e Biologia (CEDEME/UNIFESP, Brazil). Mice were group-housed (3-4 per cage), separated by sex, in polypropylene cages (44 × 34 × 16 cm) maintained on ventilated racks (Alesco, Brazil) under a 12 h light/dark cycle (lights on at 7:00 a.m.). Standard chow (Nuvilab CR-1, Brazil) and water were provided *ad libitum* throughout the experiment.

For slice electrophysiology and fiber photometry experiments at NIH, 8- to 10-week-old male and female Npas1-Cre-tdTm and Nr4a1-eGFP mice were housed in the National Institute on Alcohol Abuse and Alcoholism (NIAAA) animal facility on a reverse 12-h light cycle (lights off at 7:00 a.m.). Mice were group-housed (3-4 per cage), separated by sex, in polypropylene cages (44 × 34 × 16 cm) maintained on ventilated racks under a 12-h light/dark cycle (lights on at 7:00 a.m.). Standard chow (NIH: NIH-31 open formula diet) and water were provided *ad libitum* throughout the experiment.

Animals were allowed 7 days of acclimation to the vivarium before experimental procedures and were regularly handled by the experimenter during this period. Environmental enrichment was provided in the form of cardboard paper rolls, and cages were cleaned and replaced weekly.

### Viral vectors

Cre-dependent adeno-associated viral (AAV from Addgene) vectors were used for opto-evoked patch-clamp recordings, chemogenetic manipulation and calcium imaging of GPe^NPAS1^ neurons.

For electrophysiology, we used a Cre-dependent Channelrhodopsin-2 (ChR2) strategy to selectively stimulate NPAS1 presynaptic terminals in the striatum. NPAS1 mice were bred with Nr4a1-eGFP mice to enable visualization of matrix and striosome (patch) subcompartments during recordings. The excitatory opsin vector AAV-EF1a-double floxed-hChR2(H134R)-EYFP-WPRE-HGHpA (plasmid #20298) was obtained from Addgene. This vector expresses ChR2 under the elongation factor-1α (EF1α) promoter in a Cre-dependent (double-floxed inverted open reading frame, DIO) configuration, ensuring selective expression in Npas1-Cre–positive neurons.

For chemogenetic experiments, we used Designer Receptors Exclusively Activated by Designer Drugs (DREADDs) to inhibit and stimulate Npas1 neurons. The Gi/o-coupled DREADD vector AAV8-hSyn-DIO-hM4D(Gi)-mCherry (plasmid #44362) and the Gq-coupled DREADD vector AAV8-hSyn-DIO-hM3D(Gq)-mCherry (plasmid #44361) were obtained from Addgene. Both vectors express designer receptors under the human synapsin (hSyn) promoter in a Cre-dependent (double-floxed inverted open reading frame; DIO) configuration, ensuring selective expression in Npas1-Cre-positive neurons. Control (Cre-negative) littermates received identical viral vectors and surgical procedures to control for nonspecific viral expression and ligand effects as all animals received C21 prior to EPM behaviors.

For *in vivo* calcium imaging experiments, the genetically encoded calcium indicator GCaMP8f was expressed using AAV9-CAG-FLEX-jGCaMP8f-WPRE (plasmid #162382). This vector drives robust, Cre-dependent expression of GCaMP8f under the CAG promoter. Prior to injection, the viral preparation was diluted to a final titer of 2.3 × 10¹² viral genomes per milliliter (vg/mL). All viral vectors were delivered bilaterally into the GPe as described below.

### Stereotaxic surgery

Npas1-Cre-tdTm mice (13-15 weeks) underwent stereotaxic surgery to allow infusion and Cre-dependent expression of inhibitory and excitatory DREADDs or GCaMP8f. In Brazil, mice were weighed and received an anti-inflammatory injection (meloxicam, 0.5 mg/kg, s.c.; União Química, Brazil) before being anesthetized with inhaled isoflurane (induction: 3-5%; maintenance: 1-2%; Syntec, Brazil). The head region was cleaned and sterilized before animals were positioned in a stereotaxic apparatus (Kopf Instruments, USA). Sterile saline (0.9%) was applied to the eyes to prevent dryness, and the incision site was disinfected with povidone-iodine. Viral vectors were bilaterally injected into the GPe at the following coordinates relative to bregma (in mm): AP -0.4, ML ±2.0, DV -4.0 from dura. Vectors were delivered using a sharp glass pipette (pulled with a B100-30-7.5HP puller, Sutter Instruments, USA) in a volume of 100 nL per site. The pipette was left in place for 10 min before slow withdrawal, and the incision was closed using 6-0 nylon monofilament sutures (Shalon, Brazil). After surgery, animals were kept on a heated pad until recovery from anesthesia and received postoperative anti-inflammatory treatment (meloxicam, 0.5 mg/kg, s.c.) for three days.

In the USA, Npas1-Cre-tdTm mice (bred or not with the Nr4a1-eGFP) underwent stereotaxic surgery at the National Institutes of Health (NIH) for Cre-dependent expression of ChR2, for slice electrophysiology experiments, or GCaMP8f and implantation of optical fibers for terminal photometry recordings. Mice were anesthetized with inhaled isoflurane delivered via a Kent Scientific digital vaporizer, using 3.5% isoflurane for induction and 1.2-1.6% for maintenance at a 50 mL/min flow rate. Animals were positioned in a three-axis ultra-precision stereotaxic frame (RWD Life Science), and ophthalmic ointment was applied to prevent corneal drying. The incision site was disinfected prior to surgery. AAV9-CAG-FLEX-jGCaMP8f-WPRE was bilaterally injected into the GPe at the following coordinates relative to bregma (in mm): AP -0.4, ML ±2.0, DV -4.0 from dura mater, using the same targeting strategy as for chemogenetic experiments. Viral injections were performed using a Hamilton Neuros syringe fitted with a beveled needle (12° tip angle) at a rate of 25 nL/min, for a total volume of 100 nL per site. Following completion of each injection, the needle was left in place for 6 minutes to allow for diffusion, followed by a 4-minute automated withdrawal to minimize backflow along the injection tract.

### Confocal Imaging

Sections prepared for slice electrophysiology from Npas1-Cre-tdTm/Nr4a1-eGFP mice were fixed in 4% paraformaldehyde overnight, transferred to PBS and mounted in Vectashield (Vector Laboratories, Burlingame, CA) and imaged using a Zeiss LSM 510 Meta confocal scan head mounted on a Zeiss Axio Observer Z1 inverted microscope (Zeiss, Oberkochen, Germany). Because the tissue expressed endogenous fluorophores (eGFP and tdTomato), no immunolabeling or secondary amplification was performed. To accommodate increased scattering in thicker sections and to preserve signal integrity, excitation and emission settings were optimized for each native fluorophore: eGFP was imaged using the FITC filter set (excitation 450-490 nm, dichroic 495 nm, emission 500–550 nm), and tdTomato using the Rhodamine filter set (excitation 532–558 nm, dichroic 565 nm, emission 570–640 nm). Images were collected with Zeiss Plan-Apochromat 20×/0.8 DIC II and C-Apochromat 63×/1.2 W DIC III water-immersion objectives (Immersol W, Zeiss). To capture the large-scale organization of striosome (GFP⁺) and matrix (GFP⁻) compartments, we acquired tile-scan mosaics of the dorsal striatum using automated stitching, providing panoramic reconstructions of compartment boundaries across the mediolateral axis. For compartment-specific analyses and fine morphological inspection, z-stack images (2 µm optical sections, 2 µm pinhole) were collected through the full thickness of the fluorescent region. Laser power and detector gain were adjusted to avoid pixel saturation and kept consistent across animals processed in parallel.

### Slice electrophysiology

Npas1-Cre-tdTm/Nr4a1-eGFP mice were deeply anesthetized with isoflurane and decapitated after confirming loss of reflexes. The brain was rapidly removed and transferred to ice-cold, oxygenated sucrose-based cutting solution containing (in mM): 194 sucrose, 30 NaCl, 4.5 KCl, 26 NaHCO₃, 1.2 NaH₂PO₄, 10 glucose, 1 MgCl₂, saturated with 95% O₂ / 5% CO₂, following the slicing procedures described^45^. Coronal slices (250 μm) encompassing the dorsal striatum were cut on a vibratome and transferred to a holding chamber containing ACSF (in mM): 124 NaCl, 4.5 KCl, 26 NaHCO₃, 1.2 NaH₂PO₄, 10 glucose, 1 MgCl₂, 2 CaCl₂, equilibrated with 95% O₂ / 5% CO₂. Slices recovered for 30-60 min at 32°C and were subsequently maintained at room temperature until recording. For whole-cell voltage-clamp recordings, hemislices were transferred to a recording chamber superfused with ACSF at 30-32°C (flow rate ∼2 ml/min). Neurons in the dorsal striatum were visualized with an upright microscope equipped with epifluorescence to identify matrix and striosomal compartments (GFP^+^ vs GFP^−^). Patch electrodes (2-4 MΩ) were made from 1.5 mm OD/1.12 mm ID borosilicate glass with a filament (Worlds Precision Instruments, Sarasota, FL) pulled with a P-97 Sutter Instruments (Novato, CA) puller and filled with a high-chloride internal solution (in mM): 150 CsCl, 10 HEPES, 2 MgCl₂, 0.3 Na-GTP, 3 Mg-ATP, 0.2 BAPTA-4Cs, 5 QX-314, adjusted to pH 7.3 and 290-295 mOsm. This configuration allowed isolation of GABA-A-mediated currents as inward IPSCs at negative holding potentials. Neurons were visualized using an upright microscope (Scientifica, Uckfield, East Sussex, UK) with a LUMPlanFL N × 40/0.80 W objective (Olympus, Waltham, MA). Recordings were obtained using a Multiclamp 700A amplifier, Digidata 1322A digitizer and analyzed using pClamp 10.3 software (Molecular Devices, Sunnyvale, CA). A low-pass filter of 2 kHz and sampling frequency of 10 kHz were used. Cells were voltage-clamped at -70 mV. AMPA and NMDA receptors were blocked by bath application of 10 μM DNQX disodium salt (Cat. 2312, Tocris Bioscience) and 50 μM DL-AP5 sodium salt (ab120271, Abcam, Cambridge, MA). Optogenetic stimulation was achieved using 470-nm light pulses (2 ms, 0.05-0.9 mW) delivered through an X-Cite fluorescence illumination system equipped with a 470-nm excitation filter set. For compartment-specific comparisons, recordings were alternated between matrix (GFP^−^) and striosome (GFP^+^) neurons within the same slice, keeping light intensity and internal and external solutions identical across conditions.

### Optical fiber implantation

Following GCaMP8f viral delivery in the GPe, optical fibers were implanted bilaterally above the dorsal striatum to enable recording of GPe^NPAS1^ terminal activity. Fiber implants consisted of 200-µm core diameter optical fibers with a numerical aperture of 0.37, housed in 3-mm-length metal ferrules (Neurophotometrics). Fibers were implanted at the following coordinates relative to bregma (in mm): AP +0.5, ML ±2.0, DV −2.75 from dura, targeting the dorsal striatum bilaterally.

Optical fibers were secured to the skull using OptiBond Universal Etchant followed by Tetric EvoFlow blue-light-curable dental cement, and protected with plastic dummy ferrule caps between recording sessions. Optical fiber implantation was performed during the same surgical procedure as viral injection. Mice were allowed to recover for five weeks prior to behavioral testing and photometry recordings. Fiber placement was confirmed post hoc by histological verification of ferrule tracks and GCaMP expression within the dorsal striatum. No animals were excluded due to misplaced fiber implants.

### Elevated plus maze

Five weeks after stereotaxic surgery, mice were evaluated in the elevated plus maze (EPM). The apparatus consisted of a wooden maze elevated 50 cm above the floor, comprising two open arms and two closed arms of equal length (27.8 cm). Closed arms were enclosed by 14 cm-high walls, and all arms converged on a central square platform (7.8 × 7.8 cm). Light intensity was approximately 46 lux in the open arms and 16 lux in the closed arms. The maze was coated with waterproof varnish to prevent urine absorption.

Experiments were conducted during the light phase of the light/dark cycle (between 12:00 and 3:00 p.m.). Prior to testing, animals were acclimated to the experimental room for 30 min. The DREADD agonist Compound 21 (C21; 1.0 mg/kg, i.p.; ≥98% purity; Sigma-Aldrich, #SML2392) was administered 15 min before behavioral testing to all animals, including Cre-negative controls. At the start of each session, mice were placed in the center of the maze facing an open arm and allowed to freely explore the apparatus for 10 min. Each animal was tested only once and returned to its home cage immediately after the session. Behavior was recorded using a webcam (Logitech C920, 30 fps) connected to a computer in an adjacent room. The apparatus was cleaned with a 20% ethanol solution between sessions.

### Behavioral sequence segmentation

To characterize the structure and sequencing of exploratory behavior during EPM exploration, continuous behavior was segmented into discrete exploratory sequences based on spatial position and manually annotated risk assessment behaviors. Video recordings were first processed using SqueakPose Studio to extract frame-wise measures of body position, kinematics, posture, and head orientation. These pose-derived variables were used for behavioral quantification but were not used to automatically classify behavioral sequences.

Behavioral events and sequence boundaries were manually annotated using BORIS based on video inspection, informed by pose and trajectory information. Annotated events included sequence initiation, center crossing, risk assessment onset, and sequence termination. Risk assessment behaviors were defined by stereotyped head-out postures, body stretches, and edge dips directed toward the open arms.

Exploratory sequences were initiated when animals occupied the distal portion of an arm (sequence start zone) and terminated upon crossing ≥50% of the length of a subsequent arm. Based on the presence and outcome of intermediate risk assessment behaviors, sequences were classified into four categories: closed-to-closed transitions (C→C), NoGo sequences, Go sequences, and closed-to-open transitions (C→O). Sequence-specific timestamps were used for subsequent behavioral quantification and event-aligned photometry analyses.

### Fiber photometry recordings

Fiber photometry recordings were performed using a Neurophotometrics FP3001 system. Excitation light at 470 nm (GCaMP-dependent signal) and 415 nm (isosbestic control) was delivered through the same optical fiber using a time-multiplexed illumination scheme. Fluorescence emission was collected through the implanted fibers and detected using the FP3001 photoreceiver.

Signals were acquired at 40 Hz, synchronized to the photometry camera acquisition. Excitation light power was set to 50 µW at the fiber tip for each wavelength, measured prior to recording at the patch-cable interface. Photometry signals were recorded continuously throughout the EPM session.

### Photometry preprocessing

Raw photometry signals were processed using custom analysis scripts in Python as previously described^46^. The 470 nm and 415 nm fluorescence signals were first low-pass filtered using a zero-phase Butterworth filter with a 10-Hz cutoff, applied at the native acquisition sampling rate to filter system noise.

To correct for motion-related and shared artifacts, the filtered 415 nm isosbestic signal was linearly scaled to the 470 nm signal using a RANSAC regression approach. The scaled 415 nm signal was subtracted from the 470 nm signal to yield a motion-corrected fluorescence trace. Each recording trace and RANSAC fit were visually inspected prior to computing dF/F from the fit.

To further reduce high-frequency noise, dF/F traces were smoothed using a one-dimensional Gaussian filter (σ = 4 samples), applied independently to each peri-event trace defined by animal, hemisphere, and event number. dF/F traces were z-scored within animal across the whole EPM session and the resulting smoothed signal zdF/F was used for all subsequent analyses.

### Event alignment and time binning

Normalized and smoothed zdF/F traces were aligned to behaviorally defined event timestamps. For risk assessment sequences (Go and NoGo), signals were aligned to the onset of the first risk assessment behavior. For closed-to-closed transitions, signals were aligned to the time of center crossing. Event-aligned traces were extracted across a ±3-second peri-event window.

Peri-event traces were binned into 1/15-second time bins prior to downstream analyses. Each peri-event trace was treated as an individual observational unit defined by animal × hemisphere × event number, unless otherwise specified.

### Generalized estimating equation (GEE) modeling

Sequence-dependent temporal dynamics of GPe^NPAS1^ terminal activity were analyzed using a generalized estimating equation (GEE) framework based on previously described work^47^. The dependent variable was zdF/F, and models were fit at the level of individual peri-event traces. Time was modeled using cubic B-spline basis functions (df = 4), and behavioral sequence type was included as a categorical predictor. Sequence-specific temporal profiles were captured by interacting sequence type with the spline time terms.

Models were fit using a Gaussian family with identity link function and an autoregressive correlation structure of order 1 [AR(1)] to account for within-trace temporal dependence. Robust (sandwich) standard errors were used for inference. Overall and pairwise differences in temporal profiles were assessed using joint Wald χ² tests.

### Bootstrap procedures

Bootstrap resampling was used to estimate uncertainty around population-level photometry signals and model-predicted time courses based on previously described work^48^. Animals were resampled with replacement at the level of individual mice, and all associated peri-event traces were included for each resampled animal. Bootstrap distributions were generated using 1,000 resamples. Ninety-five percent confidence intervals were used to visualize uncertainty around population-average trajectories, and 99% confidence intervals were used for time-resolved significance visualization.

### Statistical analysis

Statistical analyses were performed using Python 3.12 with NumPy, SciPy, pandas, statsmodels, patsy, pingouin, pycircstat2, and matplotlib. Classical statistical tests, including t-tests and one-and two-way mixed ANOVAs, were conducted using pingouin, with false discovery rate (FDR) correction applied for post hoc comparisons. Distributional comparisons used Kolmogorov-Smirnov tests, and circular statistics used Rayleigh tests.

All tests were two-sided, assumed normality, and used a significance threshold of α = 0.05. No data points were excluded as outliers, and animals were randomly assigned to experimental groups. Data are reported as mean ± SEM unless otherwise specified. Exact p-values and test statistics (t, F, χ²) are reported in figure legends where applicable.

### Grants

- Coordenação de Aperfeiçoamento de Pessoal de Nível Superior (CAPES): 88887.695072/2022-00 (BDS)
- 88887.917418/2023-00 (KPA)
- Fundação de Amparo à Pesquisa do Estado de São Paulo (FAPESP): 2019/01686-0 and 2019/22009-6 (KPA)
- NIAAA, NIH 1ZIAAA000407-22 (DML)
- PRAT fellowship program of NIGMS, NIH 1FI2GM154674-02 (DLH)
- NIAAA, NIH AA025991 (AGS)

## Acknowledgements

“This research was supported by the Intramural Research Program of the National Institutes of Health (NIH). The contributions of the NIH author(s) are considered Works of the United States Government. The findings and conclusions presented in this paper are those of the author(s) and do not necessarily reflect the views of the NIH or the U.S. Department of Health and Human Services.”

## Notes

### Competing Interest Statement

The authors have declared no competing interest.

