## Supplemental Data Figure Legends for "External Globus Pallidus Arkypallidal Circuit Dynamics Gate Risk-Taking Behavior"

**Supplementary Data 1 - Representative Closed-to-Closed transition in the elevated plus maze.**
Representative overhead video of a mouse performing a Closed-to-Closed transition in the elevated plus maze. Colored pose-estimation keypoints are overlaid on the animal throughout the sequence. Video is shown at real speed (30 frames/s). The clip illustrates movement from one closed arm, through the center, and into the opposite closed arm without open-arm commitment.

**Supplementary Data 2 - Representative NoGo sequence in the elevated plus maze.**
Representative overhead video of a mouse performing a NoGo sequence in the elevated plus maze. Colored pose-estimation keypoints are overlaid on the animal throughout the sequence. Video is shown at real speed (30 frames/s). The clip illustrates approach toward the center and open-arm junction without subsequent commitment to open-arm exploration.

**Supplementary Data 3 - Representative Go sequence in the elevated plus maze.**
Representative overhead video of a mouse performing a Go sequence in the elevated plus maze. Colored pose-estimation keypoints are overlaid on the animal throughout the sequence. Video is shown at real speed (30 frames/s). The clip illustrates risk assessment followed by forward commitment into the open arm.

**Supplementary Data 4 - Representative Closed-to-Open transition in the elevated plus maze.**
Representative overhead video of a mouse performing a Closed-to-Open transition in the elevated plus maze. Colored pose-estimation keypoints are overlaid on the animal throughout the sequence. Video is shown at real speed (30 frames/s). The clip illustrates traversal through the center followed by commitment to an open arm.
